# Long-lasting structural and functional maturation in transplanted human neurons reprogrammed from glial progenitor cells

**DOI:** 10.64898/2025.12.02.691915

**Authors:** Efrain Cepeda-Prado, Christina A. Stamouli, Gianluigi Nocera, Constanza Aretio-Medina, Srisaiyini Kidnapillai, Andreas Bruzelius, Jessica Giacomoni, Janko Kajtez, Malin Parmar, Daniella Rylander Ottosson

## Abstract

Direct reprogramming of brain-resident glial progenitor cells (GPCs) into induced neurons (iN) shows great promise for future regenerative therapies for brain diseases. This type of cell conversion is facilitated by introducing lineage-specific transcription factors or small molecules to somatic cells, and can even be achieve *in vivo*, targeting resident glia cells in the brain for *in vivo* conversion. While this have shown promising results in animal models, a crucial step is to show the long-term survival and maturation of human reprogrammed cells *in vivo.* However, this remains largely unexplored partly due to the inaccessible cell source of GPC and difficulty in transplanting converted neurons. In this study, we assessed human GPCs derived iNs for their long-term structural and functional neuronal maturation after transplantation to the immunodeficient mouse brain. GPCs were transduced with established transcription factor cocktail and transplanted to the medial prefrontal cortex as three-dimensional cultures to improve survival and integration upon transplantation. Our results reveal survival of hiNs, for up to 10 months post-transplantation (MPT) and wide distribution across local and proximal functionally connected brain regions. Morphological analysis revealed a gradual neuronal maturation over time, characterized by increase in cellular complexity and expression of neuronal markers. Supporting these findings, electrophysiological recordings demonstrated progressive functional maturation up to 5 MPT with increase in sodium and potassium currents, and the ability to generate multiple evoked and spontaneous action potentials, validating a neuronal phenotype. Importantly, these changes were absent in control grafts of non-converted hGPCs that retained their glia identity. Our findings indicate that hGPCs derived iNs maintain a stable neuronal phenotype that persist in the mouse brain for extended periods. These cells exhibit survival and remarkable adaptability to the host environment, enabling both structural and functional maturation over time. These results highlight their potential as a promising strategy for neural replacement therapies aimed at restoring damaged circuits and promoting brain repair.

**Highlights:** • Human neurons reprogrammed from glia progenitor cells survive long term upon transplantation in the immunodeficient mouse brain.

• The human induced neurons migrate to local and distal connected brain regions within prefrontal cortex circuitry.

• Grafted neurons gradually develop neuronal complex morphology and electrophysiological functional properties in the mouse brain.

## Introduction

Regeneration of brain circuits after neurological injury or diseases through neuronal replacement is becoming an achievable goal, driven by rapid advancements in stem cell-based technologies (1). Among the strategies emerging in this field, direct cell reprogramming, also known as lineage conversion, shows particular promise. This reprogramming approach is termed "direct" because it bypasses a pluripotent intermediate stage, enabling the conversion of somatic cells, such as glial cells, into induced neurons (iNs) (2). Cell conversion is accomplished by overexpressing lineage-specific transcription factors, often referred to as "master regulators," that define neuronal identity. Importantly, this method enables the *in vivo* targeting of resident brain cells, e.g., glia cells, that can be converted into neurons with minimal risk of tumor formation (3).

Recent *in* reprogramming strategies in mouse models have demonstrated successful reprogramming of resident glial cells into functional neurons (4–7) that can contribute to circuit repair (8, 9). Among different glial populations, glial progenitor cells (GPCs; also known as oligodendrocyte progenitor cells or NG2 cells) are particularly well-suited for reprogramming due to their connectivity with neuronal networks, ability to proliferate in adulthood, and their widespread distribution throughout the brain (10). However, applying these strategies to hGPCs has been challenging because their late embryonic developmental limits the access to primary human tissue cultures (11).

To address this limitation, we utilize a differentiation protocol to derive hGPCs from embryonic stem cells (12, 13). These GPC can be reprogrammed into functional iNs in two-dimensional cultures through the ectopic expression of specific neuronal transcription factors (12, 14). Recently, we adapted our reprogramming protocols to a three-dimensional (3D) culture environment, which provides a more physiologically relevant microenvironment that enhances the structural integrity and improves survival of hiNs upon transplantation into the adult rodent brain (15). With this approach, we obtained functional human iNs *in vitro* that exhibit a transcriptional profile comparable to *bona fide* interneurons (16). However, the survival migration and functional maturation of hiNs *in vivo* remain largely unexplored (17).

In this study, we aimed to investigate the long-term survival and adaptability of hGPCs derived hiNs after transplantation to the immunodeficient mouse brain. Human GPCs were first reprogrammed into iNs in 3D spheroids and implanted into the medial prefrontal cortex, where they were allowed to mature over a period of 1, 3, 5 and 10 months post transplantation (MPT). Our findings show that hiNs survived for up to 10 MPT in the mouse brain with remarkable migration across various brain regions. The hINs exhibited progressive structural and functional maturation that became evident after three months by dynamic change in ion channel activity and the development of complex dendritic arbors. Taken together, these findings demonstrate, for the first time, a long-term survival and gradual maturation of grafted hiNs generated from GPCs and highlight the potential of reprogramming strategies in the development of future neuronal replacement therapies.

## Methods

### Cell line and culturing conditions

GPCs differentiation started from a human embryonic stem cell (hESC) line known as RC17 (Roslin Cells Ltd, cat.no. hPSCreg RCe021-A, see (18)), as previously described (12, 13). Briefly, hESCs were cultured on LN521-coated plates with (0.5 mg/cm^2^, Biolamina, Sundbyberg, Sweden) and kept in IPS-Brew XF medium (StemMACS, Miltenyi Biotec, Bergisch Gladbach, Germany). Weekly passages were performed using EDTA (0.5 mM, Gibco, Thermo Fisher, Waltham, MA, USA). Human ESC-derived s were maintained in glial medium (GM) containing DMEM/F12 basal medium, B27 supplement, N1 supplement (Sigma-Aldrich), MEM NEAA, Antibiotic-Antimycotic, T3 (60 ng/ml, Sigma-Aldrich), db-cAMP (1 μM, SigmaAldrich), Biotin (100 ng/ml, Sigma-Aldrich), recombinant human PDGF-AA protein (10 ng/ml, R&D Systems), recombinant human IGF-I (10 ng/ml, R&D Systems) and recombinant human NT-3 Protein (10 ng/ml, R&D Systems). Fluorescence-activated cell sorting (FACS) analysis was used to estimate the proportion of oligodendrocyte biased progenitors (CD140a^+^/CD44^−^), astrocyte biased (CD140a^−^/CD44^+^) and a bipotential (CD140a^+^/CD44^+^) progenitors within the hESC-derived GPC population as described in (16).

### Direct reprogramming of 3D hGPCs cultures into induced neurons

Direct conversion of hGPCs to neurons was initiated by overexpression five transcription factors, three constitutively expressed (*Ascl1*, *DLX5* and *LHX6*) under control of the phosphoglycerate kinase (PGK) promoter for ubiquitous expression, and two doxycycline-inducible system (*Sox2* and *FOXG1*) driven by doxycycline inducible promoter (Tet-On system) and co-transduced with a TET-ON trans-activator (#20342, Addgene, Watertown, MA, USA). Third-generation lentiviral vectors (LVs) produced as previously described (19) and mixed with the cells during seeding to ensure homogenous exposure (14). For *in vivo* tracking, cells were co-transduced with a lentivirus expressing green fluorescent protein (GFP) under the human Synapsin 1 promoter (pLV-hSyn1-eGFP; Addgene #177810, Watertown, MA, USA) enabling identification of transplanted human cells during histological analysis and electrophysiological recordings. Viral titers ranged from 2.5×10^8^ to 3.42×10^9^ transducing units as estimated by qPCR.

After 2-3 weeks from thawing, hGPCs were scraped and dissociated into single cells suspension using Accutase (StemPro, Thermo Fisher; Waltham, MA, USA). Human GPCs were mixed with the lentiviral reprogramming factors with a multiplicity of infection of 1–2 per vector and approximately 1.2x10^6^ cells were seeded in each well of a 24-well microplate, where each well contains approximately 300 microwells with 800 μm in size (AggreWell™800, STEMCELL Technologies Cambridge, UK). This microwell arrays promotes spheroid formation through self-aggregation. After 24 hours, fresh medium containing doxycycline (5 mg/ml, Duchefa, Haarlem, Netherlands) was added to induce Tet-On expression. Three days later, medium was replaced with neural differentiation (NDiff) medium, which consisted of NDiff227 (Takara-Clontech, Gothenburg, Sweden), supplemented with doxycycline (5 mg/ml), small molecules (CHIR99021, 2 mM, Axon, Groningen, Netherlands); SB-431542 (10 mM, Axon, Groningen, Netherlands); noggin (0.5 mg/ml, R&D Systems R&D Systems, Minneapolis, MN, USA); LDN193189 (0.5 mM, Axon, Groningen, Netherlands); VPA (1 mM, Merck Millipore, Burlington, MA, USA) and growth factors (LM-22A4, 2 mM, R&D Systems, Minneapolis, MN, USA); GDNF,(2 ng/ml, R&D Systems, Minneapolis, MN, USA); NT3,10 ng/ml, R&D Systems, Minneapolis, MN, USA); db-cAMP (0.5 mM, Sigma-Aldrich, St. Louis,MO, USA). NDiff medium was refreshed every 2-3 days. Small molecules and doxycycline were maintained for 12 DIV, after which the spheroids were used for transplantation.

A subset of both non-converted and converted spheroids was used to verify glia and pan neuronal markers expression by immunofluorescent and RT-qPCR to confirm activation of viral *Ascl1* (a pioneer proneuronal transcription factor), transgene using specific primers.

### Animals and cell transplantation

Immunodeficient NSG mice (NOD.Cg-Prkdc scid Il2rg_tm1Wjl / SzJ; Jackson Laboratory, strain 005557,) were used on postnatal day 14 (P14). All experimental procedures were conducted in accordance with the European Union Directive 2010/63/EU on the protection of animals used for scientific purposes and were approved by the Ethical Committee for Animal Research at Lund University and the Swedish Department of Agriculture (permit no. 13978-18 and 14836/23)

For cell transplantation a stereotactic surgery was carried out under depth anesthesia with 4% induction, 1-2% maintenance) in a mixture of air and nitrous oxide (N_2_O) at a 4:1 ratio. Anesthetized animals were carefully placed in a stereotaxic frame. Analgesia (0.1 mg/kg of Buprenorphine) was administered subcutaneously before the surgery started. Quartz glass capillaries (0.8 - 1mm tip) were used to inject the spheroids in the mPFC (coordinates A/P: +2.3 and M/L: -0.7 from bregma) in two deposits (D/V: -1.0 and -1.5) on the right hemisphere. Each deposit, 0.5 μL with approximately 75.000 cells/ μL was injected at a rate of 0.2μL/min with a 2 min diffusion time. Post-surgery the animals were continuously monitored, maintained under standard conditions (water and food *ad libitum,* 12 h light:12 h dark cycle). Data was collected from a total of 28 mice of both sexes.

### Immunocytochemistry

Animals were sacrificed at 1, 3, 5 and 10 MPT. Briefly, animals were anesthetized with an intraperitoneal overdose of sodium pentobarbital (60 mg/kg) and perfused intracardially with room-temperature (RT) 0.9% saline for 1min followed by ice-cold 4% PFA (pH 7.4 ± 0.2) for 8 minutes. The brains dissected and post-fixed overnight in 4% PFA at 4°C and then cryoprotected in 25% sucrose for 24h. Brains were sectioned at 35 µm thickness using a tissue slicing Microtome (Leica SM2010 R) for histological analyses. Additionally, acute slices (200–275 µm thickness) prepared for electrophysiological recordings were also used for histology and neuronal reconstructions.

Immunostainings were carried out on free floating brain sections in 12-well plates (VWR, ref. 734-2799). Sections were washed three times in 0.1M potassium phosphate buffer saline (3XKPBS) and incubated with citrate buffer, pH 6.0 (Sigma, Ref: C9999) for 45 min at 60°C for antigen retrieval, followed by 3X-KPBS washes. For 3, 3’-diaminobenzidine (DAB) stainings, sections were incubated in a quenching solution (10% H_2_O_2_ and 10% methanol in KPBS) for 15 min to block endogenous peroxidase activity, washed 3X-KPBS, and then incubated for 1h in blocking solution (0.3% Triton X-100 in KPBS, containing 5% donkey serum) at RT. Next, primary antibody (mouse anti-hNCAM, Santa Cruz, sc106, 1:1000) were prepared in the blocking solution and left on a shaker at RT overnight. The following day the sections were washed 3X-KPBS and preincubated with blocking solution at RT for 30min followed by 1hour in secondary antibodies (horse Anti-Mouse IgG Antibody, Biotinylated, Vectorslab, BA 2001, 1:200). An avidin-biotin complex (ABC Complex) was used after the secondary antibody for 1h and washed 3X-KPBS. The sections were then incubated in 0.05% DAB-solution for 2min before adding 20μL 3% H_2_O_2_ to allow color development. Finally, the sections were washed 3XKPBS, mounted onto chrome alum-gelatin coated slides, dehydrated in an ascending gradient of alcohols, cleared in xylene and cover slipped with DPX mounting media.

For immunofluorescence, stainings were carried out for both spheroids and tissue sections. The samples were incubating for 1 hour in blocking solution followed by 24 hours incubation with primary antibodies prepared in blocking solution. For this study the following primary antibodies were used: chicken anti GFP (Abcam, AB13970, 1:1000), mouse anti hTAU-HT7 (Thermo, MN 1000, 1:300), mouse anti human nuclear antigen (HuNu, Millipore, MAB 128, 1:200) and goat anti human PDGFRa (R&D Systems, AF-307-NA, 1:300). Fluorophore-conjugated secondary antibodies (Alexa-488, Alexa-568, or Alexa-647, 1:200; Jackson ImmunoResearch Laboratories, West Grove, PA, USA) in combination with DAPI (1:2000, Sigma-Aldrich, St. Louis, MO, USA) were applied for 1h at RT on a shaker. The tissue sections were washed 3X-KPBS, mounted onto Fisherbrand Superfrost Plus slides (fisher scientific, 12-550-15) and cover slipped with PVA-DABCO mounting medium. Spheroids were directly transferred to 96-well plates with flat and clear bottoms (Ibidi, Gräfelfing, Germany) for confocal microscopy.

### RNA extraction, cDNA synthesis, RT-qPCR

Spheroids were lysed in RLT buffer (Qiagen, Hilden, Germany), and total RNA was extracted using the RNeasy Micro Kit (Qiagen, Hilden, Germany) according to the manufacturer’s instructions. RNA was reverse transcribed to cDNA with the Maxima First Strand cDNA Synthesis Kit (Thermo Fisher, Waltham, MA, USA). RT-qPCR reactions were prepared using the Bravo Automated Liquid Handling Platform (Agilent, Santa Clara, CA, USA), combining cDNA with SYBR Green Master Mix and gene-specific primers in technical triplicates. qPCR was run on a LightCycler 480 II using a 40-cycle, two-step protocol (95°C denaturation, 60°C annealing/extension). Relative gene expression was calculated by the ΔΔCT method, normalized to GAPDH and hGPCs prior to lentiviral transduction.

### Confocal images and morphological analysis

Confocal images from spheroids after 12DIV were acquired using 10x air objective of Zeiss 780 Confocal Laser-Scanning Inverted Microscope with Zeiss Zen Blue Edition software.

For tissue samples, images of GFP-expressing human neurons were taken at 1, 3, 5, and 10 MPT using 10× air and 40× oil immersion objectives. Images were acquired with a Leica Stellaris 5 Confocal Laser Scanning Microscope and digitized using LAS X software (Leica Microsystems). To capture high-resolution images across large fields of view, multi-panel acquisitions were conducted and automatically stitched using the microscope’s native software.

Detailed morphological analyses were carried out using IMARIS 10.2 (Oxford Instruments). Images were collected as z-stack series along the z-axis at 0.35 µm intervals to ensure full threedimensional integrity during reconstruction. Soma and dendritic arborization analysis were analyzed using the Filament Tracer tool in IMARIS, with the soma defined as the seeding point for filament reconstruction. Quantitative parameters included number of branch segments and number of terminal points.

Dendritic complexity was assessed using Sholl analysis in ImageJ/Fiji with the Neuroanatomy plugin. Filament tracing was performed using NeuronJ, from which the total number of neurites and overall neuron extension length were quantified.

### Electrophysiological recordings

Recordings were performed at 1, 3, and 5 MPT. Briefly, animals were anesthetized via intraperitoneal injection of sodium pentobarbital (60 mg/kg) and transcardially perfused with ∼10 mL of oxygenated (95% O₂ / 5% CO₂) N-methyl-D-glucamine (NMDG)-HEPES artificial cerebrospinal fluid (aCSF) containing (in mM): 92 NMDG, 2.5 KCl, 1.25 NaH₂PO₄, 30 NaHCO₃, 20 HEPES, 25 glucose, 2 thiourea, 5 sodium ascorbate, 3 sodium pyruvate, 0.5 CaCl₂·2H₂O, and 10 MgSO₄·7H₂O. The pH was adjusted to 7.3–7.4 using 37% HCl (7 ± 0.2 mL), and osmolarity was set to 300–305 mOsm/kg.

Brains were rapidly removed and placed in ice-cold NMDG-HEPES aCSF. Coronal acute slices (275 µm) containing the mPFC were prepared using a vibratome (Leica VT1200 S, Germany). Slices were incubated for 25 minutes at 35 °C, then transferred to a room temperature holding chamber containing HEPES-aCSF containing (containing (in mM): 124 NaCl, 2.5 KCl, 1.25 NaH₂PO₄, 24 NaHCO₃, 12.5 glucose, 5 HEPES, 2 CaCl₂·2H₂O, and 2 MgSO₄·7H₂O (pH 7.4, 300–305 mOsm/kg), until recordings commenced.

GFP-expressing transplanted cells were visualized using a 40× water-immersion objective on a fixed-stage Olympus BX51WI microscope equipped with an IR-CCD camera and LED illumination system (Prizmatix Ltd., Israel). Whole-cell patch-clamp recordings were performed at room temperature under constant perfusion (1 mL/min) with oxygenated HEPES-aCSF. Patch electrodes (5–7 MΩ) were pulled from borosilicate glass using a Narishige PC-100 vertical pipette puller and filled with an intracellular solution containing (in mM): 130 K-gluconate, 10 KCl, 0.2 EGTA, 10 HEPES, 4 Mg-ATP, 0.5 Na-GTP, and 10 Na-phosphocreatine (pH 7.25, 296 mOsm/kg). Recordings were acquired using a Multiclamp 700B amplifier and pClamp 11 software (Molecular Devices, USA). Signals were low pass filtered at 0.1 kHz and digitized at 2 kHz. Passive membrane properties, including resting membrane potential (RMP), capacitance (Cm), were obtained directly from pClamp, data acquisition software, at the start of each recording. Series resistance was compensated by bridge balance.

Recordings were performed at a holding potential of −70 mV, and cells requiring a holding current >100 pA were excluded. Action potentials (APs) were evoked by injecting current steps ranging from −20 to +35 pA (500 ms duration, 5 pA increments) and by ramping the cells up form 0 to 50pA over the course of 500ms. AP threshold (APt), amplitude (APh) and afterhyperpolarization (AHP) were calculated from first AP triggered in response to increase depolarizing current. Voltage-gated inward sodium (Na⁺) and outward potassium (K⁺) currents were recorded under voltage-clamp conditions with 100 ms depolarizing steps from −70 mV to +40 mV in 10 mV increments. Data were analyzed using Clampfit 11 (Molecular Devices) and Igor Pro 9 (WaveMetrics, USA), with the NeuroMatic plugin (20).

### Statistics and data analysis

Data were analyzed and visualized using GraphPad Prism v10.2 (GraphPad Software, San Diego, California, USA). Sample sizes and the specific statistical tests applied are indicated in the main text and figure legends.

## Results

### Direct neuronal reprogramming of hGPCs in 3D cultures

Human GPCs were first generated from hESCs using multiple independent cell batches. Identity was confirmed by the expression of surface markers defining the different populations within the hGPCs using flow cytometry. As previously described (12, 13), hGPC population was mainly compose by oligodendrocyte biased progenitors (CD140^+^/CD44^−^), followed by a smaller proportion astrocyte biased (CD140^−^/CD44^+^) and a subset co-expressing both markers (Supple. Figure 1A, Table S1).

**Figure 1.**
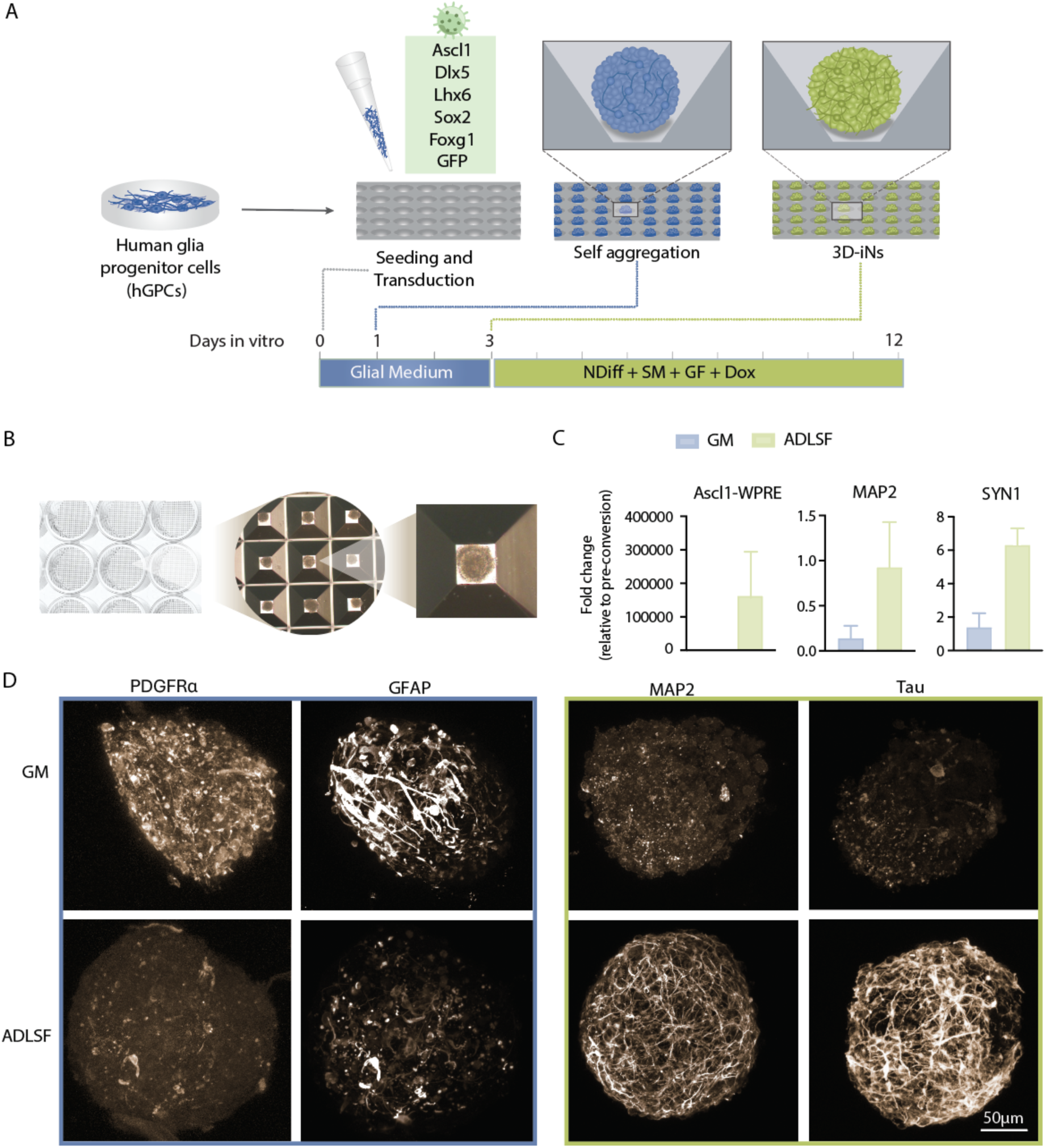
Generation of 3D-induced human neuronal microcultures. (A) The panel presents a schematic timeline of the 3D direct reprogramming protocol utilized to generate hiNs from hGPCs. (B) Brightfield images illustrate 24-well ultra-low attachment plates, along with a close-up of the microwells containing spheroids at 7 days in vitro (DIV). (C) The real-time quantitative RT-qPCR (RT-qPCR) analysis compares the expression of lentiviral introduced *Ascl1* gene, and the pan-neuronal markers known as microtubule-associated protein 2 (*MAP2*) and Synapsin 1 (*SYN1*) in both hGPCs and hiNs following reprogramming. (D) Confocal images from immunostaining shows the differential expression of glial markers (GFAP, PDGFRα) and neuronal markers (TAU, MAP2) before (indicated by a light blue border) and after reprogramming (indicated by a light green border). Data represent results from three independent experiments conducted at 12 DIV and are presented as mean ± SEM in C.

After hGPCs expansion in 2D cultures, 3D cultures were generated by detaching the cells through mechanical dissociation and seeded into 24-well ultra-low attachment plates to promote uniform spheroid formation (Figure B). A combination of five transcription factors (TFs) were used for conversion, *FOXG1*, Sox2, *Ascl1*, *DLX5*, and *LHX6*, defined as ADLSF and delivered via an independent lentiviral vectors (16). To enable tracking of the cells after transplantation, we co-delivered a lentiviral vector encoding green fluorescent protein (GFP). The cells were maintained in glial medium (GM) for 3 DIV to facilitate efficient viral transduction before transitioning into neurodifferentiation medium (NDiff) supplemented with small molecules (SMs) and doxycycline (DOX) to induce Sox2 and Foxg1 expression. Under these conditions, converted and non-converted spheroids (hGPCs maintained in GM) were maintained for an additional 12 DIV (Figure 1A).

To confirm a neuronal conversion, we evaluated gene and protein expression prior to transplantation. Real-Time Quantitative PCR (RT-qPCR) showed upregulation of the lentiviral-introduced *Ascl1* gene, which is a crucial proneuronal TF. Additionally, there was an increased expression of pan-neuronal genes, including synapsin I (*SYN1*) and microtubule-associated protein 2 (*MAP2*), as illustrated in Figure 1C. Consistent with this, immunostaining of ADLSF transfected cells showed increased expression of neuronal markers microtubule-associated protein 2 (MAP2) and TAU (Figure 1E). These markers were absent in non-converted hGPCs that instead showed high levels of pan-glial markers, platelet-derived growth factor receptor alpha (PDGFRα) and glial fibrillary acidic protein (GFAP). These results confirm that ADLSF efficiently reprograms hGPCs into hiNs in a 3D environment, consistent with previous findings (14, 16).

### Long term survival and extensive spatial distribution of hiNs in the mouse brain

To assess the ability of hGPCs-derived iNs to adapt and integrate into the host mouse brain, spheroids were stereotactically injected into the secondary motor cortex (M2) within the mPFC of NSG immunodeficient mice at postnatal day 14 (P14). Survival and spatial distribution of grafted cells were assessed at 1-, 3-, 5-, and 10 MPT (Figure 2A).

**Figure 2.**
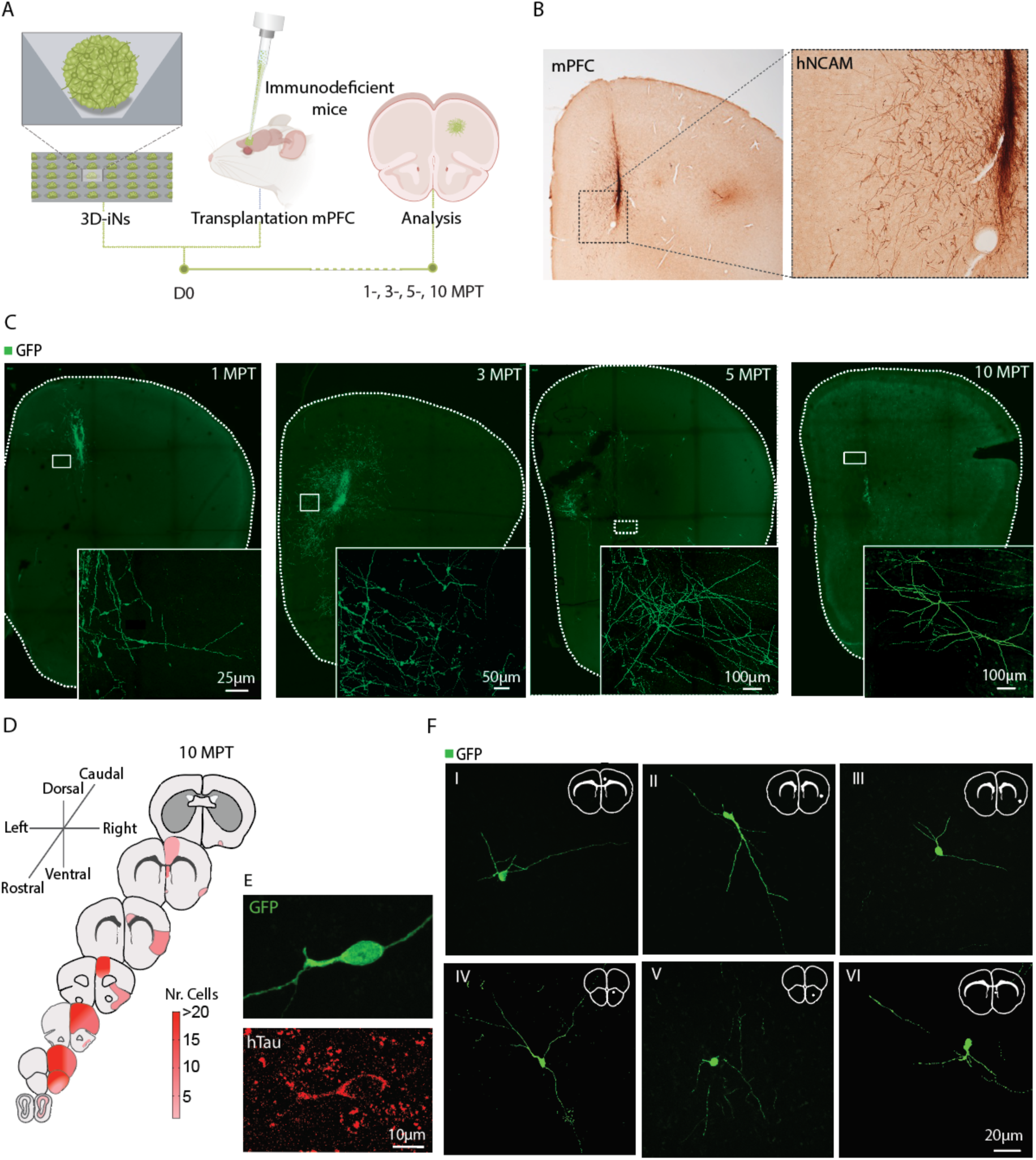
**Transplantation of 3D-microcultures and long-lasting survival of hiNs in the mouse brain.** (A) The illustration depicts the intracerebral injection of 3D microcultures into the mouse brain, along with the defined experimental endpoints relative to the day of transplantation (D0). These endpoints include assessments at 1-, 3-, 5-, and 10 MPT. (B) Diaminobenzidine (DAB) staining shows the location of the grafted hiNs and confirms their survival at 1 MPT. (C) Fluorescence confocal images of the graft core demonstrate the prolonged survival of hiNs and developing complex dendritic arbors over time. (D) Schematic representation of mouse brain slices, ranging from rostral to caudal, illustrating the spatial distribution of transplanted cells. The number of cells in each region is indicated by a color-coded bar. (E) Confocal image of a grafted human cell at 10 MPT expressing human-specific Tau protein, which is typically present in mature neurons. (F) Fluorescence confocal images display representative images of individual hiNs found in various mouse brain regions including secondary motor cortex (I), claustrum (II), piriform cortex (III), the taenia tecta of the olfactory bulb (IV), the granular layer of the olfactory bulb (V) and the septum (VI) at 10 MPT.

Cell survival was first confirmed at 1 MPT through immunohistochemical staining using the human-specific neural cell adhesion molecule (hNCAM) antibody. At this stage, hNCAMpositive cells were primarily localized in the mPFC and had started infiltrating the mouse brain parenchyma (Figure 2B). Additional immunofluorescence analysis of GFP-positive cells further confirmed a widespread distribution of hiNs from the injection side (Figure 2C). At 1 MPT, the cells displayed an immature bipolar morphology and were predominantly located in the graft core (close-up in Figure 2C), which remained distinguishable until 3 MPT; but was no longer detectable at 5 and 10 MPT, likely due to extensive cell migration to other brain regions (Figure 2D, F) and partial cell loss. Nevertheless, the remaining hiNs exhibited a continuous development with characteristic neuronal features, and prominent projections with numerous ramifications (see Figure 2C).

By 10 MPT, the transplanted cells showed extensive neuronal structure, increased size, and a remarkable distribution throughout the host brain. Human iNs were found across the mouse neocortex and in several other regions, particularly those that were functionally connected to the mPFC circuit (21, 22) (Figure 2D). Furthermore, they were positive for human-specific Tau (hTau), a microtubule-associated protein, confirming the neuronal phenotype (Figure 2E). Representative images showing morphological characteristics of hiNs located in distinct brain regions, including the secondary motor cortex (I), claustrum (II), piriform cortex (III), the taenia tecta of the olfactory bulb (IV), the granular layer of the olfactory bulb (V) and the septum (VI) (Figure 2F), revealing their capabilities to integrate into the mouse brain parenchyma.

This study is the first to demonstrate that hiNs can survive up to 10 MPT with wide distribution over long distances in an *in vivo* environment.

### Human iNs undergo continuous structural neuronal maturation for up to 10 MPT in the mouse brain

After confirming the survival and widespread distribution of hiNs, we next examine their morphological changes over time. Immunostaining for hGFP revealed a characteristic neuronal morphology in animals transplanted with ADLSF-transduced cells (Figure 3A). This was strikingly different from the oligodendrocyte-like morphology, seen in control animals (transplanted with GPF-transduced GPCs) characterized by small somas and multiple fine, branched processes (Figure 3A, Suppl. Figure 1B). A crucial first step in the structural analysis was to confirm the human identity of GFP-positive hiNs using human-specific nuclear antigen (HuNu) histological assessment. The results demonstrated a clear co-localization of GFP and HuNu for ADLSF-transduced GPCs (Figure 3B). Similarly, all non-converted hGPCs exhibited co-expression of HuNu (see Suppl. Figure 1C). Importantly, no endogenous mouse cells expressed GFP expression in any samples.

**Figure 3.**
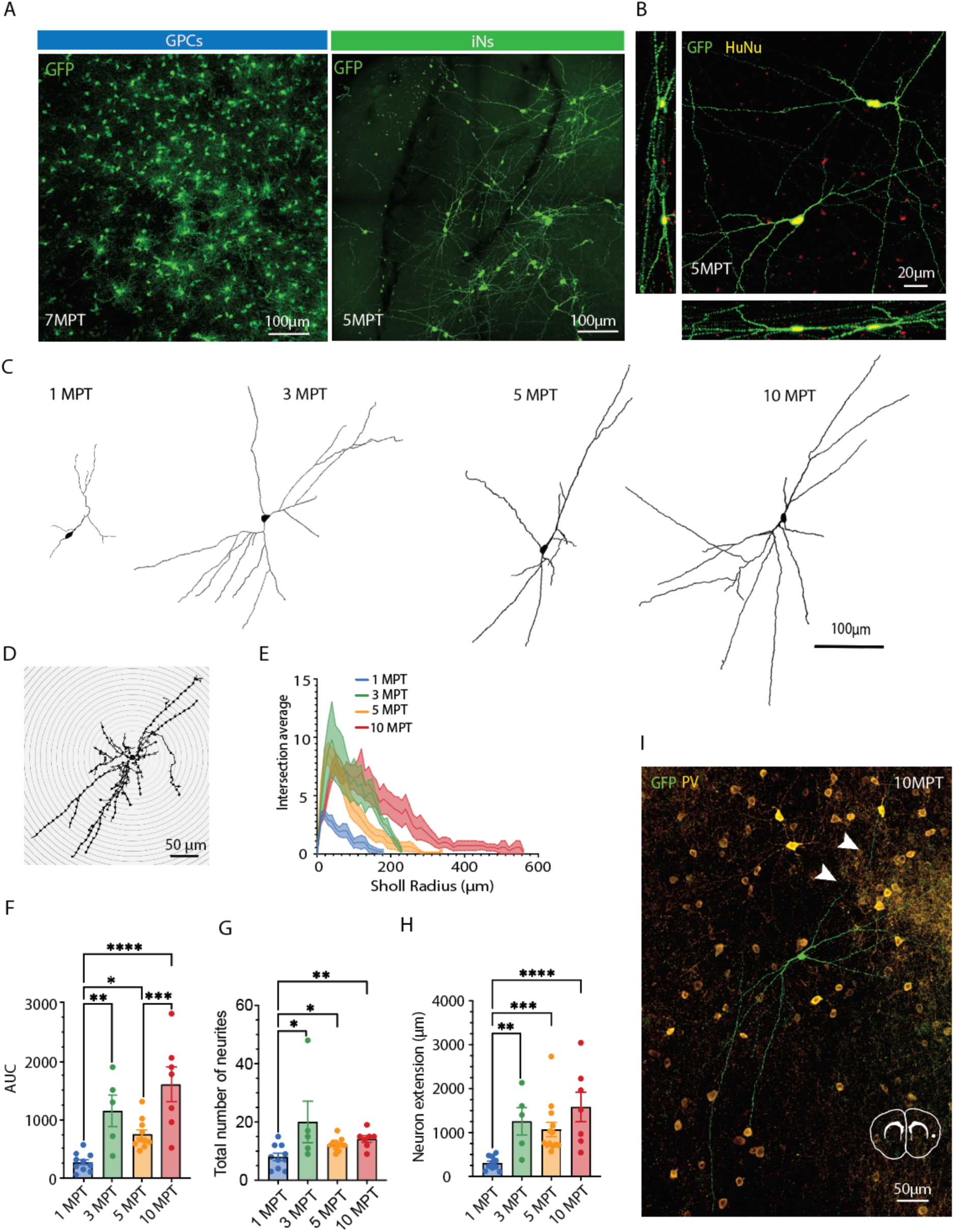
**Structural characterization of hiNs following transplantation.** (A) Representative confocal images from mPFC illustrating the overall morphological modifications from GPCs to neuron conversion. (B) A maximum intensity projection of a fluorescence confocal image stack illustrates the overlap of green fluorescent protein (GFP)positive hiNs with a human-specific nuclear marker (HuNu) in the medial prefrontal cortex (mPFC). The inserts provide an orthogonal view confirming the colocalization of the two markers. (C) 3D reconstructions of grafted hiNs demonstrate the time-dependent structural maturation. (D) The inset provides an example of Sholl analysis, while (E) presents a Sholl plot that illustrates the degree of cellular complexity of grafted hiNs at different time points. (F) Areaunder-the-curve (AUC) of Sholl plot was calculated to assess overall branching density. (G) Quantitative analysis of number of neurites and (H) neuronal extension based maximum number of crossed concentric rings. (I) Immunofluorescent showing mouse parvalbumin (PV) positive interneurons surrounded a human GFP positive cell located in the insular cortex and projecting (white arrowheads) to another brain region. The structural data were collected from three different animals at each time point, except for 10 MPT, where cells were found in only one of four animals. Data are presented as mean ± SEM in E, F, G and H. ***p<0.0001, **p<0.001, *p<0.05, using a one-way nonparametric ANOVA test, followed by Tukey multiple comparation.

To further assess the development of neuronal morphology over time, brain samples from alltime points were stained for GFP and cell exhibiting neurotypical characteristics, including long processes and multiple branches, were selected for 3D reconstruction and quantitative analysis. Only cells that had migrated away from the graft core were included, as the dense packing within the core hindered reliable reconstruction. On average, 6 to 11 hiNs were reconstructed per time point (Suppl. Figure 2). A 3D visualization demonstrated progressive structural maturation of the hiNs at 1, 3, 5 and 10 MPT, characterized by more complex neuronal architecture (Figure 3C).

To quantify neuronal complexity, we further performed Sholl analysis (23), by drawing concentric rings around the soma at 10 µm intervals spanning a radius of 0–600 µm (Figure 3D). In this analysis, complexity was defined as the average number of neurite intersections per ring, whereas neuronal extension was quantified by the number of rings crossed by neurites (i.e., Sholl radius). We also estimated the total number of neurites per neuron.

The data show that converted cells exhibited progressive structural remodeling, resulting in distinct Sholl profiles over time (Figure 3E). At 1 MPT, most hiNs displayed a small, bipolar morphology with minimal branching, resembling immature glial cells (Figure 3C, E). By 3 MPT, neurons developed a larger and more elaborate cytoarchitecture characterized by multiple neurites and increased branching complexity. While 5 MPT, complexity modestly decreased relative to 3 MPT (Figure 3C), as indicated by fewer intersections (Figure 3E) and reduced branching density in the area-under-the-curve (AUC) analysis, yet by 10 MPT, overall complexity returned to levels comparable to those at 3 MPT (Figure 3F).

Similarly, the number of neurites per cell increased significantly from 1 MPT (8 ± 1.2) to 3 MPT (20 ± 7.5) and then stabilized (5 MPT: 14.3 ± 1.4, 10 MPT: 14.4 ± 1.2, Figure 3G). In contrast, neurite length continued to increase over time (Figure 3H), suggesting a developmental shift toward elongation of existing projections rather than generation of additional branches (Figure 3I). This is exemplified in Figure 3I, where an isolated hiN in the insular cortex at 10 MPT extends long-range projections (white arrowheads) beyond the boundaries of the tissue section, a frequent observation at this stage.

Collectively, these findings demonstrate that hiNs gradually mature into structurally complex neurons cells between 1 and 10 MPT across various mouse brain regions. While neuronal complexity is already achieved by 3-MPT, later time points promote elongated neurites rather than higher complexity with potential to connect with other transplanted human cells (see Suppl. Video).

### Gradual development of functional neuronal properties for hiNs within the mouse brain

To investigate the functional maturation of hiNs, we performed patch-clamp electrophysiological recordings at all experimental end time points specifically assessing capacitance and resting membrane potential (RMP), expression of activated Na^+^ and K^+^ channels and the ability to generate spontaneous and evoked action potentials (APs) (Figure 4A). Additionally, functional changes were also accessed in grafted hGPCs at 7.5 MP, as these cells have been reported to develop some electrophysiological properties (24). Human cells were identified based on their GFP expression and selected by their neurotypical morphology (Figure 4A).

**Figure 4.**
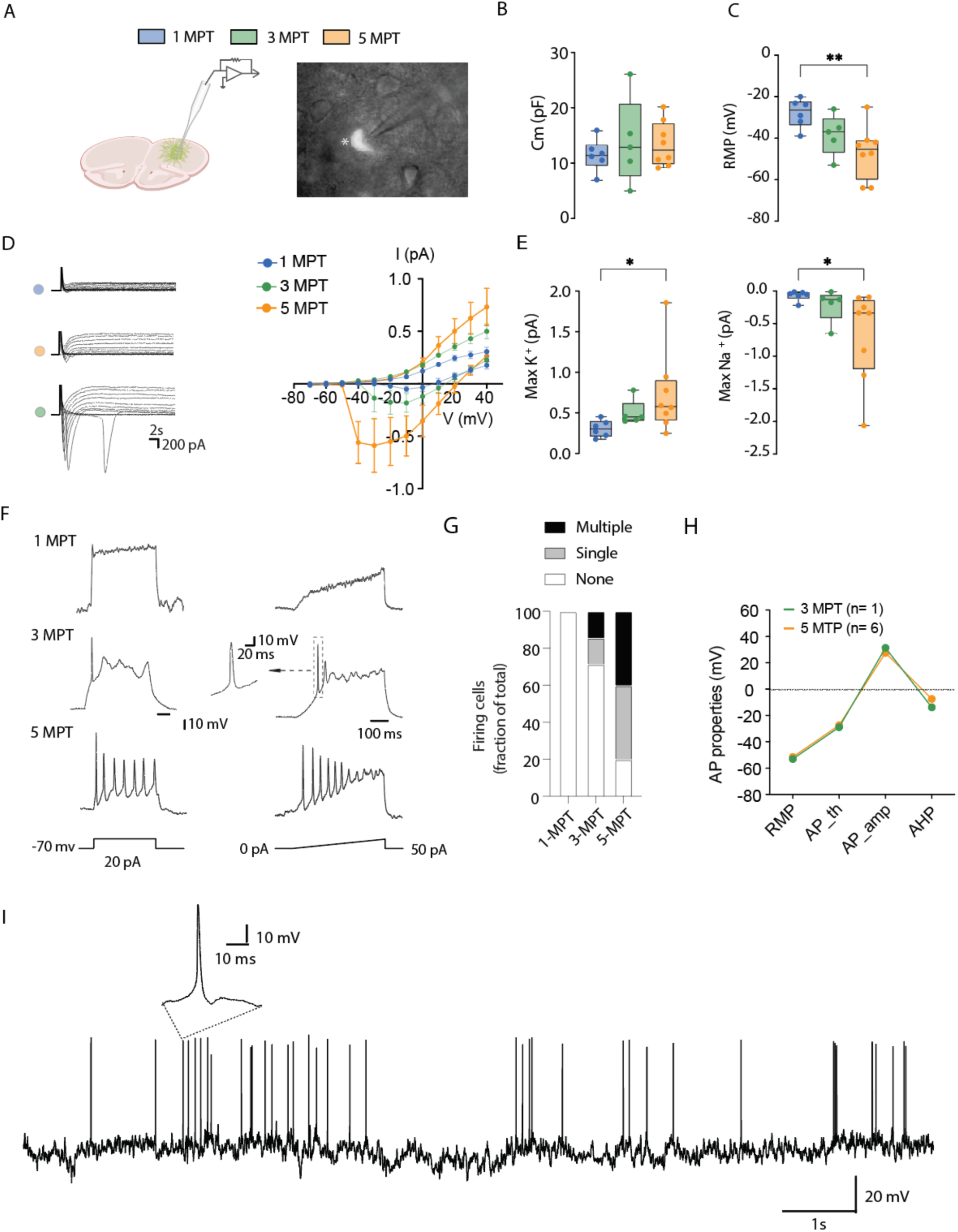
**Functional maturation of grafted hiNs.** (A) Diagram of the whole-cell patch-clamp recording setup. The right panel shows a brightfield image of a GFP-positive hiN undergoing recording (white asterisk). Passive membrane properties of human cells including (B) membrane capacitance (Cm), and (C) resting membrane potential (RMP). (D) Representative traces of inward sodium (Na⁺) and outward potassium (K⁺) currents recorded in response to depolarizing voltage steps. On the left panel, current–voltage (I–V) relationships showing mean amplitudes of Na⁺ and K⁺ currents. (E) quantification of maximum peak of inward Na⁺ and outward K⁺ current amplitudes (F) Representative traces of evoked action potentials (APs) in response to ether current injection (on the right panel) or a ramp of 50 pA of depolarization (on the left panel). (G) Proportion of active (firing) cells across different timepoints post-transplantation. (H) Quantitative analysis of evoked AP properties, including resting membrane potential (RMP), AP threshold (AP_th), AP peak amplitude (AP_amp) and afterhyperpolarization (AHP). (I) Representative trace of spontaneous action potentials recorded from grafted hiNs. Recordings were obtained from 2 -3 independent experiments. 1MPT (n=6 from 3 animals), 3MPT (n=5 from 2 animals) and 5MPT (n=8 from 3 animals). Median, interquartile ranges (IQR), and whiskers (min and max) are shown in B, C and E. Data are presented as mean ± SEM in D and F. ** p < 0.001, * p < 0.05 using a one-way nonparametric ANOVA test, followed by Dunn’s multiple comparison test.

All recorded cells were located in the mPFC, close to the graft core. The data revealed a nonsignificant change in capacitance over time (Figure 4B; median: 1MPT, -0.05 pF, 3MPT, -0.13 pF, 5MPT, -0.34 pF), suggesting no substantial alterations in soma size. However, significant differences in RMP were observed at 5MPT, with a median of -45.4mV, compared to -26.45mV at 1MPT (Figure 4C), indicating a progressive depolarization of the cell membrane, and an increased number and activity of leak channels and sodium-potassium pumps, which are essential for maintaining the voltage difference across the cell membrane at rest.

Supporting the data above, grafted hiNs exhibited a gradual increase in peak amplitude Na+ and K+ currents in response to depolarizing voltage steps, with significant changes at 5 MPT compared to 1 MPT (Figure 4D, E). These alterations could be attributed to the incorporation of more voltage-gated ion channels in the cell membrane, enhancing the likelihood of evoking APs at 3 and 5 MPT compared to 1 MPT (Figure 4F, G), and overall, the AP properties (i.e. AP threshold, AP amplitude and afterhyperpolarization (AHP) (Figure 4H). Notably, spontaneous APs were observed in two of the eight recorded cells at 5 MPT (Figure 4I).

Grafted hGPCs in control animals on the other hand, exhibited higher cell capacitance compared to hiNs and demonstrated a more negative RMP, (Suppl. Figure 1D) While they also showed some Na^+^ and K^+^ currents (Suppl. Figure 1E) as in previously reports (24) the hGPCs were only able to elicit single APs but never multiple (Supple. Figure 1F).

Altogether, these findings demonstrate that only hiNs gradually develop both passive and active membrane properties associated with neuronal function in an *in vivo* environment from 3 to 5 MPT.

## Discussion

Recent advancements in stem cell transplantation have led to significant progress with ongoing efforts exploring various cell products and sources including neuronal progenitors in clinical trial (25). Traditionally, research has focused on cell transplantation; however, promising alternatives include *in vivo* reprogramming strategies, aimed for in brain neuronal repair reducing the risks of rejection and tumor formation by utilizing the patient’s own resident cells (3)

GPCs are particularly suitable for *in vivo* reprogramming due to their ability to proliferate throughout adulthood, capacity for synaptic connectivity, and their widespread distribution throughout the adult central nervous system (26). Thus, targeting resident hGPC for patientspecific neuronal reprogramming could be an attractive alternative to cell transplantation (27). Although glial reprogramming *in vivo* has been successfully tested in rodent models targeting cortical or striatal glial cells (5, 8, 28), its application to human systems has primarily been restricted to *in vitro* models in which cells are maintained in an artificial environment (3, 17). Therefore, xenotransplantation of human cells into animal models is crucial for further investigating the long-term maturation and functional integration of directly reprogrammed human neurons in the brain (15, 29, 30). However, transplanting hiNs into animals poses challenges, including fragility and poor survival after transplantation of mature cells into a host brain (15, 31, 32). To overcome this limitation, we have introduced spheroids for transplantation rather than a traditional single-cell suspension. One significant advantage of 3D cultures, which grow in suspension, is that they eliminate the need for enzymatic or mechanical dissociation steps that compromise cell viability and disrupt intercellular interactions (15, 33). Notably, the spheroids used in this study have a small diameter (approx. 80 µm), allowing them to be injected directly into the mouse brain using standard stereotaxic surgical protocols with increased survival and integration compared to single cell suspension (15). Using this protocol we demonstrated extended survival timeframe for these hiNs from 1 up to 10 MPT, which has not been previously reported and only showed by a limited number involving stem cell-derived neurons (34–37). Moreover, the long-term characterization of the converted hiN showed development of a neuronal identity from 3 MPT to 10 MPT with increased dendritic length and neurite complexity as well as expression of essential neuronal markers, that was not seen for non-converted GPCs, confirming a transcription factor-dependent neuronal transformation.

We chose to transplant the cells the PFC because of its significant role in various diseases related neurodevelopmental disorders (38–40). This brain region is responsible for integrating signals from the present to predict the future and adapt behavior by processing information from multiple brain regions. Interestingly, the migratory patterns of the transplanted hiNs aligned with those of the functional brain regions associated with medial prefrontal circuitry (22). This alignment suggests that developmental factors likely influenced these specific migration patterns, as the neurons were transplanted at a postnatal stage when neuronal networks were still maturing. Similar findings have been observed with human cortical neurons derived from ESCs, transplanted into the lateral ventricle at postnatal age, and they successfully integrated into migratory streams, across cortical layers, and differentiated into various subtypes of cortical neurons (36).

One advantage of direct reprogramming, compared with stem-cell-based differentiation, is the rapid acquisition of key features of the target cell identity. In ADLSF-transduced GPCs, Gene Ontology enrichment analysis revealed a significant upregulation of neuronal genes involved in cytoskeletal remodeling, membrane protein activation, and localized protein synthesis as early as 13 days post-conversion, coinciding with the development of neuronal functional properties (16). Following transplantation, these cells slowly integrated into the mouse cellular microenvironment and exhibited both evoked and spontaneous multiple action potentials at 5 MPT (see Figure F–I). Despite this functional maturation, the cells remained in a juvenile state, likely reflecting species-specific differences in neuronal developmental timelines between human grafted and mouse host neurons (36). Approaches to further promote maturation may include transplanting cells at an earlier reprogramming stage (41), enhancing neuronal plasticity through exercise (42) or applying electrical stimulation (43).

On the other hand, whereas only a few hiNs persisted in the graft core up to 10 MPT compared to the earlier time points the surviving cells exhibited extensive migration across functionally connected brain regions (Figure 1), which is an important step for reconstructing neuronal networks. as even a small number of well-connected human cells could significantly influence brain function. For instance, transplantation studies have indicated that approximately 7% to 10% of the initial cell dose may be sufficient to reestablish neuronal circuits. (34). While the functional integration of hiNs and their potential to restore damaged brain circuits remains to be fully tested, this study marks a significant advancement to the field by demonstrating long term survival and response to appropriately signals from the host brain, allowing hiNs to migrate and progressively mature both structurally and functionally. These findings represent a significant step forward in developing cell repair strategies by using direct reprogramming technologies.

## Data availability

Data supporting the findings of this study are available upon reasonable request.

## Acknowledgments

This work was supported by Stem cell center and MultiPark at Lund University and funded by the Swedish Research Council [2021-01839 and 2021-03149; DRO], Knut and Alice

Wallenberg Foundation [2021-0088; DRO], Olle Engkvist Foundation [213-0229; DRO], Royal Physiographic Society in Lund [45610; ECP]. The funders had no role in study design, data collection and analysis, decision to publish, or preparation of the manuscript.

We would like to thank you Anna Hammarberg for her assistance with FACS, Emanuella

Monni for support with confocal imaging, German Ramos Passarelo, Ulla Jarl and Bengt Mattsson for their assistance with sample preparation, histology and cell transplantation.

## Authors’ contributions

All authors approved the final version of the manuscript. Conceptualization: ECP, DRO; formal analysis: ECP, GN, CAM; funding acquisition: ECP, DRO; investigation: ECP, CS, GN, CAM, SK; project administration: ECP, SK, DRO; resources/samples: JAB, JG, MP; supervision: ECP, DRO; visualization: ECP, CS, GN, CAM, JK; writing - original draft: ECP, DRO; writing - review & editing: ECP, CS, GN, CAM, SR, JAB, JG, JK, MP, DRO.

## References

1. De Luca M, Aiuti A, Cossu G, Parmar M, Pellegrini G, Robey PG. Advances in stem cell research and therapeutic development. Nat Cell Biol. 2019;21(7):801–11.

2. Bocchi R, Masserdotti G, Gotz M. Direct neuronal reprogramming: Fast forward from new concepts toward therapeutic approaches. Neuron. 2022;110(3):366–93.

3. Barker RA, Gotz M, Parmar M. New approaches for brain repair-from rescue to reprogramming. Nature. 2018;557(7705):329–34.

4. Chouchane M, Melo de Farias AR, Moura DMS, Hilscher MM, Schroeder T, Leao RN, et al. Lineage Reprogramming of Astroglial Cells from Different Origins into Distinct Neuronal Subtypes. Stem Cell Reports. 2017;9(1):162–76.

5. Heinrich C, Gascon S, Masserdotti G, Lepier A, Sanchez R, Simon-Ebert T, et al. Generation of subtype-specific neurons from postnatal astroglia of the mouse cerebral cortex. Nat Protoc. 2011;6(2):214–28.

6. Pereira M, Birtele M, Shrigley S, Benitez JA, Hedlund E, Parmar M, et al. Direct Reprogramming of Resident NG2 Glia into Neurons with Properties of Fast-Spiking Parvalbumin-Containing Interneurons. Stem Cell Reports. 2017;9(3):742–51.

7. Torper O, Ottosson DR, Pereira M, Lau S, Cardoso T, Grealish S, et al. In Vivo Reprogramming of Striatal NG2 Glia into Functional Neurons that Integrate into Local Host Circuitry. Cell Rep. 2015;12(3):474–81.

8. Lentini C, d’Orange M, Marichal N, Trottmann MM, Vignoles R, Foucault L, et al. Reprogramming reactive glia into interneurons reduces chronic seizure activity in a mouse model of mesial temporal lobe epilepsy. Cell Stem Cell. 2021;28(12):2104-21 e10.

9. Qian C, Dong B, Wang XY, Zhou FQ. In vivo glial trans-differentiation for neuronal replacement and functional recovery in central nervous system. FEBS J. 2021;288(16):4773–85.

10. van Tilborg E, de Theije CGM, van Hal M, Wagenaar N, de Vries LS, Benders MJ, et al. Origin and dynamics of oligodendrocytes in the developing brain: Implications for perinatal white matter injury. Glia. 2018;66(2):221–38.

11. Yeung MS, Zdunek S, Bergmann O, Bernard S, Salehpour M, Alkass K, et al. Dynamics of oligodendrocyte generation and myelination in the human brain. Cell. 2014;159(4):766–74.

12. Nolbrant S, Giacomoni J, Hoban DB, Bruzelius A, Birtele M, Chandler-Militello D, et al. Direct Reprogramming of Human Fetal- and Stem Cell-Derived Glial Progenitor Cells into Midbrain Dopaminergic Neurons. Stem Cell Reports. 2020;15(4):869–82.

13. Wang S, Bates J, Li X, Schanz S, Chandler-Militello D, Levine C, et al. Human iPSCderived oligodendrocyte progenitor cells can myelinate and rescue a mouse model of congenital hypomyelination. Cell Stem Cell. 2013;12(2):252–64.

14. Giacomoni J, Bruzelius A, Stamouli CA, Rylander Ottosson D. Direct Conversion of Human Stem Cell-Derived Glial Progenitor Cells into GABAergic Interneurons. Cells. 2020;9(11):2451.

15. Kajtez J, Laurin K, Nilsson F, Bruzelius A, Cepeda-Prado E, Birtele M, et al. Threedimensional cell-cell interactions promote direct reprogramming of patient fibroblasts into functional and transplantable neurons. Sci Adv. 2025;11(23):eadq7855.

16. Stamouli C-A, Degener A, Cepeda-Prado E, Bruzelius A, Andersson E, Giacomoni J, et al. A distinct lineage pathway drives parvalbumin chandelier cell fate in human interneuron reprogramming Sci Adv. 2025;In press.

17. Karow M, Sanchez R, Schichor C, Masserdotti G, Ortega F, Heinrich C, et al. Reprogramming of pericyte-derived cells of the adult human brain into induced neuronal cells. Cell Stem Cell. 2012;11(4):471–6.

18. De Sousa PA, Tye BJ, Bruce K, Dand P, Russell G, Collins DM, et al. Derivation of the clinical grade human embryonic stem cell line RCe021-A (RC-17). Stem Cell Res. 2016;17(1):1–5.

19. Zufferey R, Nagy D, Mandel RJ, Naldini L, Trono D. Multiply attenuated lentiviral vector achieves efficient gene delivery in vivo. Nat Biotechnol. 1997;15(9):871–5.

20. Rothman JS, Silver RA. NeuroMatic: An Integrated Open-Source Software Toolkit for Acquisition, Analysis and Simulation of Electrophysiological Data. Front Neuroinform. 2018;12:14.

21. Bhattarai JP, Etyemez S, Jaaro-Peled H, Janke E, Leon Tolosa UD, Kamiya A, et al. Olfactory modulation of the medial prefrontal cortex circuitry: Implications for social cognition. Semin Cell Dev Biol. 2022;129:31–9.

22. Le Merre P, Ahrlund-Richter S, Carlen M. The mouse prefrontal cortex: Unity in diversity. Neuron. 2021;109(12):1925–44.

23. Sholl DA. Dendritic organization in the neurons of the visual and motor cortices of the cat. J Anat. 1953;87(4):387–406.

24. Livesey MR, Magnani D, Cleary EM, Vasistha NA, James OT, Selvaraj BT, et al. Maturation and electrophysiological properties of human pluripotent stem cell-derived oligodendrocytes. Stem Cells. 2016;34(4):1040–53.

25. Kirkeby A, Main H, Carpenter M. Pluripotent stem-cell-derived therapies in clinical trial: A 2025 update. Cell Stem Cell. 2025;32(1):10–37.

26. Dimou L, Gotz M. Glial cells as progenitors and stem cells: new roles in the healthy and diseased brain. Physiol Rev. 2014;94(3):709–37.

27. Nakatsukasa Y, Yamada Y, Yamada Y. Research of in vivo reprogramming toward clinical applications in regenerative medicine: A concise review. Regen Ther. 2025;28:12–9.

28. Pereira M, Birtele M, Rylander Ottosson D. In Vivo Direct Reprogramming of Resident Glial Cells into Interneurons by Intracerebral Injection of Viral Vectors. J Vis Exp. 2019(148).

29. Zhou H, Wen P, Liu Y, Ye Z, Xiong W, Liu Y, et al. MiR-138 reprograms dental pulp stem cells into GABAergic neurons via the GATAD2B/MTA3/WNTs axis for stroke treatment. Biomaterials. 2026;325:123618.

30. Papetti AV, Jin M, Ma Z, Stillitano AC, Jiang P. Chimeric brain models: Unlocking insights into human neural development, aging, diseases, and cell therapies. Neuron. 2025;113(14):2230–50.

31. Berg LJ, Lee CK, Matsumura H, Leinhaas A, Konang R, Shaib AH, et al. Human neural stem cells directly programmed from peripheral blood show functional integration into the adult mouse brain. Stem Cell Res Ther. 2024;15(1):488.

32. Colasante G, Lignani G, Rubio A, Medrihan L, Yekhlef L, Sessa A, et al. Rapid Conversion of Fibroblasts into Functional Forebrain GABAergic Interneurons by Direct Genetic Reprogramming. Cell Stem Cell. 2015;17(6):719–34.

33. Rossen NS, Anandakumaran PN, Zur Nieden R, Lo K, Luo W, Park C, et al. Injectable Therapeutic Organoids Using Sacrificial Hydrogels. iScience. 2020;23(5):101052.

34. Bershteyn M, Broer S, Parekh M, Maury Y, Havlicek S, Kriks S, et al. Human pallial MGE-type GABAergic interneuron cell therapy for chronic focal epilepsy. Cell Stem Cell. 2023;30(10):1331–50 e11.

35. Bershteyn M, Zhou H, Fuentealba L, Chen C, Subramanyam G, Cherkowsky D, et al. Human stem cell-derived GABAergic interneuron development reveals early emergence of subtype diversity and gradual electrochemical maturation. Neuron. 2025;113(19):3162–84 e10.

36. Linaro D, Vermaercke B, Iwata R, Ramaswamy A, Libe-Philippot B, Boubakar L, et al. Xenotransplanted Human Cortical Neurons Reveal Species-Specific Development and Functional Integration into Mouse Visual Circuits. Neuron. 2019;104(5):972–86 e6.

37. Shrestha S, Anderson NC, Grabel LB, Naegele JR, Aaron GB. Development of electrophysiological and morphological properties of human embryonic stem cell-derived GABAergic interneurons at different times after transplantation into the mouse hippocampus. PLoS One. 2020;15(8):e0237426.

38. Xu P, Chen A, Li Y, Xing X, Lu H. Medial prefrontal cortex in neurological diseases. Physiol Genomics. 2019;51(9):432–42.

39. Chini M, Hanganu-Opatz IL. Prefrontal Cortex Development in Health and Disease: Lessons from Rodents and Humans. Trends Neurosci. 2021;44(3):227–40.

40. Jobson DD, Hase Y, Clarkson AN, Kalaria RN. The role of the medial prefrontal cortex in cognition, ageing and dementia. Brain Commun. 2021;3(3):fcab125.

41. Espuny-Camacho I, Michelsen KA, Gall D, Linaro D, Hasche A, Bonnefont J, et al. Pyramidal neurons derived from human pluripotent stem cells integrate efficiently into mouse brain circuits in vivo. Neuron. 2013;77(3):440–56.

42. Moriarty N, Fraser TD, Hunt CPJ, Eleftheriou G, Kauhausen JA, Thompson LH, et al. Exercise promotes the functional integration of human stem cell-derived neural grafts in a rodent model of Parkinson’s disease. Stem Cell Reports. 2025;20(5):102480.

43. Wang L, Yao Y, Xie B, Lei M, Li Y, Shi J, et al. Nanoelectrode-Mediated Extracellular Electrical Stimulation Directing Dopaminergic Neuronal Differentiation of Stem Cells for Improved Parkinson’s Disease Therapy. Adv Mater. 2025;37(6):e2409745.

