## Supplementary material for "Long-lasting structural and functional maturation in transplanted human neurons reprogrammed from glial progenitor cells"

**Supplementary Figure 1. Grafted hGPCs maintain their cell identity up to 7.5 MPT.**

(A) Percentages of oligodendrocyte biased ($CD140^{+}/CD44^{-}$), astrocyte biased ($CD140^{-}/CD44^{+}$) and bipotential ($CD140a^{+}/CD44^{+}$) progenitors detected different batches of GPCs by FACS. See Table S1 (B). Confocal image illustrating typical hGPCs morphology at 3MPT. (C) A maximum intensity projection of a fluorescence confocal image stack illustrates the overlap of green fluorescent protein (GFP)-positive hGPCs with a human-specific nuclear marker (HuNu) in the medial prefrontal cortex (mPFC) at 3MPT. (D) Passive membrane properties of hGPCs, including membrane capacitance (Cm) and resting membrane potential (RMP). (E) Current–voltage (I–V) relationships showing mean amplitudes of $Na^{+}$ and $K^{+}$ currents and quantification of peak inward $Na^{+}$ and outward $K^{+}$ current amplitudes of hGPCs. (F) Original traces of evoked action potentials (APs) in response to current injection from all recorded cells. Median, interquartile ranges (IQR), and whiskers (min and max) are shown in D and E. Data are presented as mean ± SEM in F. Recordings were obtained from four animals, and 13 cells were recorded and including in the analysis.

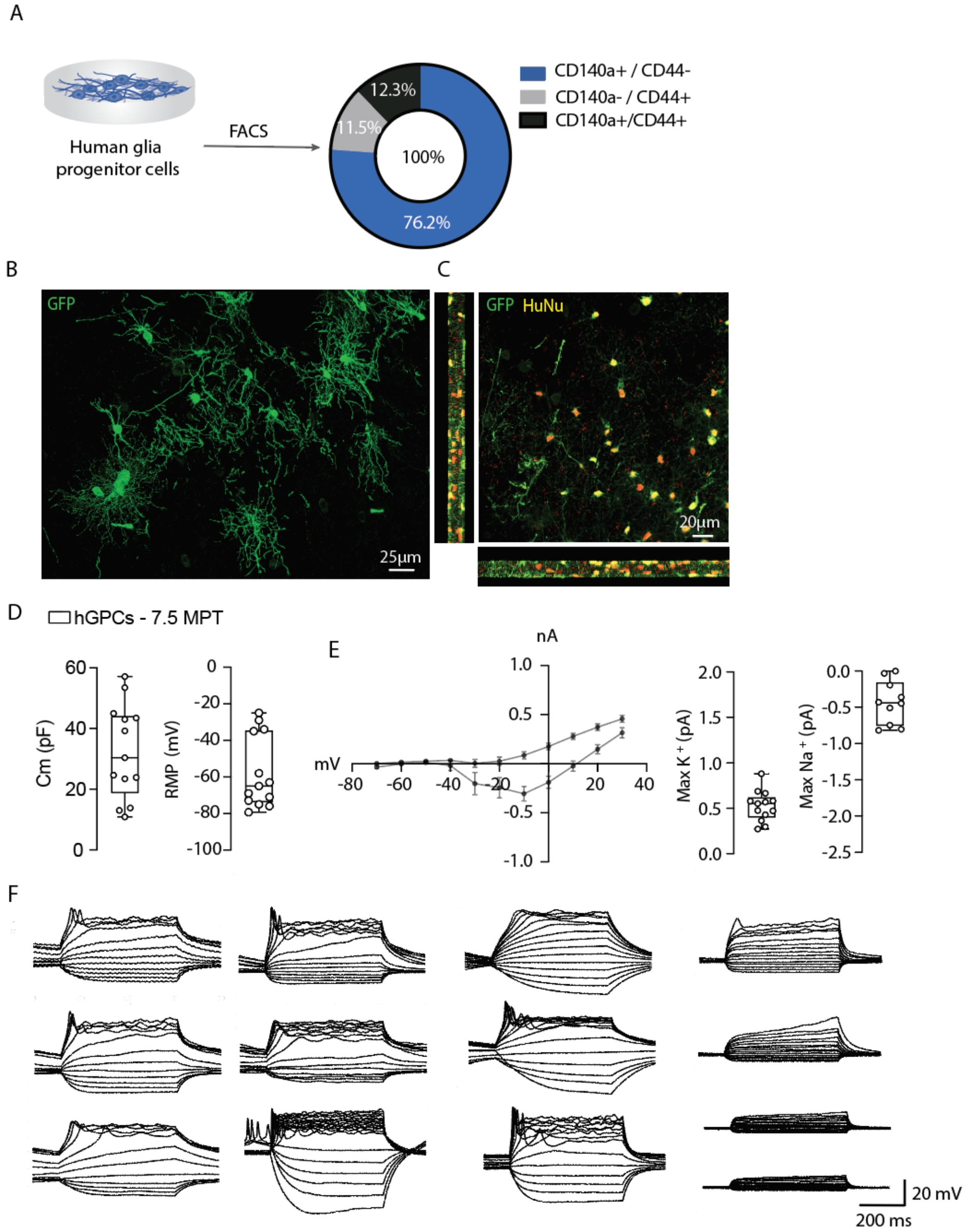

**Supplementary Figure 2. Reconstructed hiNs.** Compilation of all reconstructed cells for 1, 3, 5 and 10 months post transplantation (MPT).

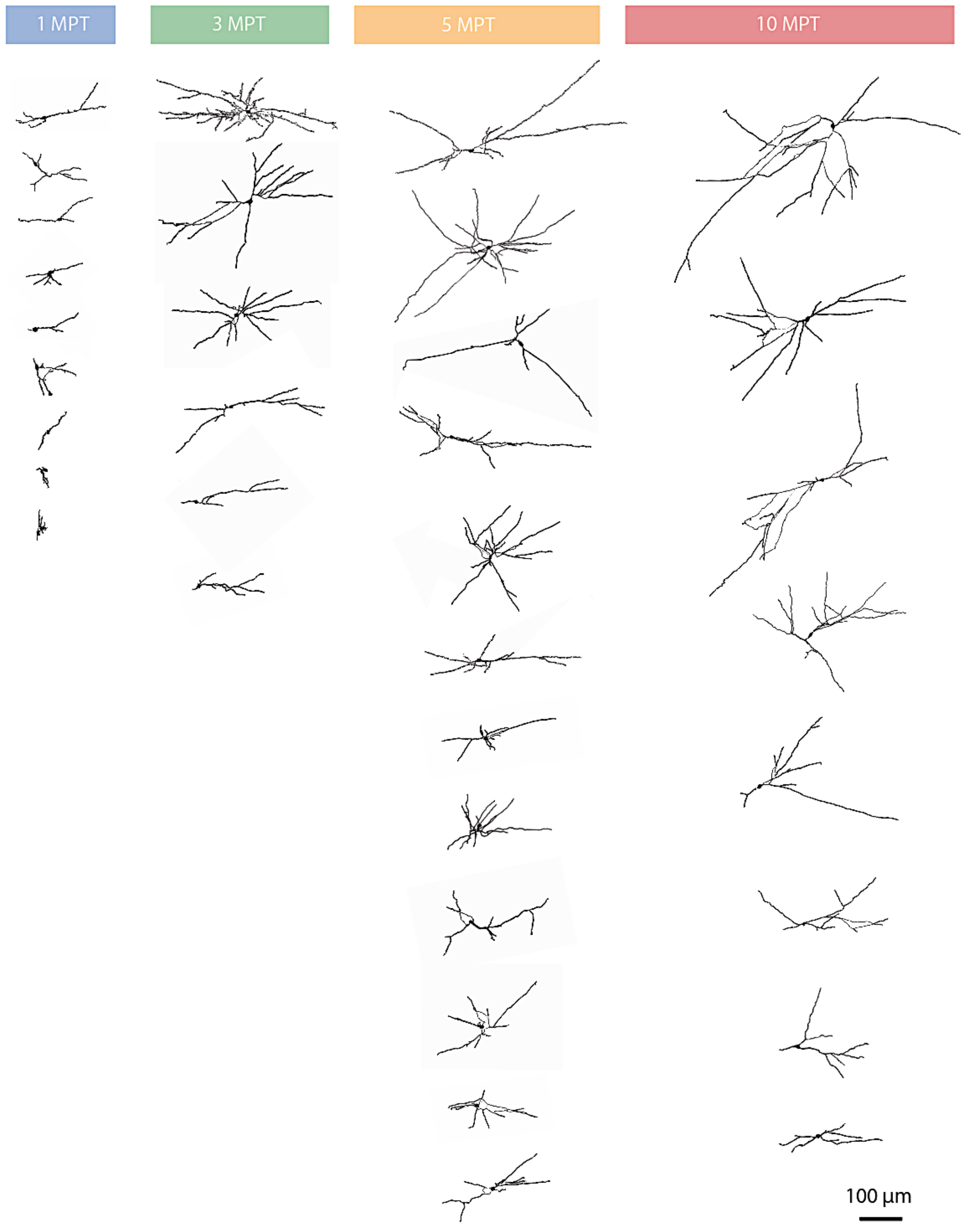

**Table S1:** Human GPCs batches used to initiate the cell reprogramming experiments. Related to supplementary Figure 1A.

| | | **Percentage of positive cells** | | |
|---|---|---|---|---|
| **hGPC batch ID** | **DIV** | **CD140a+ /CD44$^-$** | **CD140a- / CD44+** | **CD140a+/CD44+** |
| JAB01 | 221 | 56.6 | 3.4 | 11.0 |
| JAB01 | 187 | 48.2 | 22.3 | 16.7 |
| JAB05 | 206 | 70.4 | 9.6 | 5.9 |
| JAB04 | 207 | 69.6 | 1.1 | 17.2 |
| JAB04 | 208 | 74.2 | 37.5 | 13.1 |
| CAS01 | 220 | 55.4 | 0.3 | 0.3 |
| JAB05 | 172 | 68.5 | 0.5 | 11.1 |
| JAB01 | 221 | 30.9 | 6.1 | 9.8 |
| JAB01 | 221 | 47.8 | 7.3 | 6.2 |
| JAB05 | 206 | 60.7 | 0.2 | 2.8 |
| **Average** | | **58.23** | **8.83** | **9.41** |
